# Healthy aging and cognitive impairment alter EEG functional connectivity in distinct frequency bands

**DOI:** 10.1101/2023.01.24.525301

**Authors:** Wupadrasta Santosh Kumar, Supratim Ray

**Affiliations:** Centre for Neuroscience, Indian Institute of Science, Bengaluru, India, 560012, Facsimile

**Author notes:** Declaration of interests: The authors declare no competing financial interests.

**Keywords:** EEG, Gamma oscillations, healthy aging, cluster shrinkage, functional connectivity, Alzheimer’s Disease

## Abstract

Functional connectivity (FC) indicates the interdependencies between brain signals recorded from spatially distinct locations in different frequency bands, which is modulated by cognitive tasks and is known to change with aging and cognitive disorders. Recently, the power of narrow-band gamma oscillations induced by visual gratings has been shown to reduce with both healthy aging and in subjects with mild cognitive impairment (MCI). However, the impact of aging/MCI on stimulus-induced gamma FC has not been well studied. We recorded electroencephalogram (EEG) from a large cohort (N=229) of elderly subjects (>49 years) while they viewed large cartesian gratings to induce gamma oscillations and studied changes in alpha and gamma FC with healthy aging (N=218) and MCI (N=11). Surprisingly, we found that aging and disease changed power and FC in different ways. With healthy aging, alpha power did not change but FC decreased significantly. MCI reduced gamma but not alpha FC significantly compared with age and gender matched controls, even when power was matched between the two groups. Overall, our results show distinct effects of aging and disease on EEG power and FC, suggesting different mechanisms and the potential to use EEG stimulus-induced FC along with power for early diagnosis of Alzheimer’s Disease.

## Introduction

Neural coordination referred as functional connectivity (FC) (Fingelkurts et al., 2005), refers to the statistical relationship between spatially distant neurophysiological events which is generally considered to be mediated by neuronal assemblies (Mcintosh, 1999). FC can be studied at various spatial and temporal scales, using brain signals such as fMRI, electro/magneto-encephalogram (EEG/MEG) or neuronal firing activity, and is quantified using a variety of techniques such as coherence (Srinivasan et al., 2007), imaginary part of coherence (Nolte et al., 2004), phase locking value (PLV; (Lachaux et al., 2000)), pairwise phase consistency (PPC; (Vinck et al., 2010)) and so on. Many studies have reported changes in FC in different frequency bands with cognitive tasks, aging and with brain disorders (Ishii et al., 2017). For example, reduction in alpha (8-12 Hz) FC was reported with healthy aging in resting state EEG (Moezzi et al., 2019), while gamma (30-55 Hz) FC reduction was reported in autism (Safar et al., 2020), and in Alzheimer’s Disease (AD) and schizophrenia (Uhlhaas and Singer, 2006).

Majority of EEG FC studies that describe the effect of aging and cognitive disorder are during resting state, where many factors can alter the data, such as specific instructions, cognitive state before recording, caffeine intake, or random episodic spontaneous thoughts (van Diessen et al., 2015). Having a stimulus dependent paradigm restricts subject specific variations. Such responses are termed as either evoked or induced, based on their trial-wise relationship with stimulus onset (Tallon-Baudry and Bertrand, 1999). Recent studies in human EEG/MEG have shown that gamma oscillations can be induced by presentation of stationary cartesian or moving annular gratings (Orekhova et al., 2015; Murty et al., 2018; Pantazis et al., 2018), whose power changes with age (Murty et al., 2020) and mental disorders such as MCI/AD (Murty et al., 2021). However, whether aging or mental disorders change stimulus-induced gamma FC is not well studied.

Further, while both aging and cognitive disorders generally reduce the power of brain oscillations, the effect of these two factors on FC could be different. For example, Sullivan and colleagues (2019) reported that healthy aging and MCI produce distinct changes in fMRI FC (measured using bivariate-correlation) across different networks in the brain. Another study (Finnigan and Robertson, 2011) reported altered relationship between alpha (8-12 Hz) and theta (4-8 Hz) oscillations with healthy aging and pathological aging. Finding robust connectivity measures that distinguish aging versus mental disorder could help in developing better biomarkers that are more sensitive to the onset of cognitive disorders compared to power-based biomarkers.

We recorded EEG data from healthy (N=218) and MCI (N=11) subjects while they maintained fixation on large achromatic gratings presented on a monitor (same dataset as the one used in (Murty et al., 2020, 2021)). Large gratings were shown to induce two gamma oscillations, termed “slow” (20-34 Hz) and “fast” (35-65 Hz) (Murty et al., 2018; 2020, 2021). We therefore estimated FC in alpha, slow gamma and fast gamma bands using PPC, which is an unbiased estimator of the squared PLV (see (Vinck et al., 2010) for details). We first analysed the effect of aging on visual stimulus-induced FC by splitting subjects into middle-aged (50-66 yrs) and elderly (>66 yrs) groups, as done in previous studies (Murty et al., 2020, 2021). Since FC could potentially be dependent on power along with aging, we performed a power-matching procedure (see detailed below) to control for power dependence. We also tested FC dependency on power and age using a robust linear regression model. Finally, we compared the FC for MCI subjects versus their age and gender matched controls.

## Results

Figure 1 illustrates the EEG power and FC analysis for a representative subject. Figure 1a shows the topoplot of raw power in the slow gamma range between 250-750 ms after stimulus onset, while Figure 1b shows the change in power from baseline (−500 to 0 ms), revealing an increase in power in the occipital area. As in our previous studies, all analyses were performed on three groups of occipital electrodes (left, right and back; see Methods for details). Figure 1c shows the change in average power spectral density profile between stimulus and baseline periods for the left (top) and right (bottom) electrode groups (these electrodes are highlighted in Figure 1b), revealing a clear suppression in alpha and elevation in slow and fast gamma bands. To estimate FC, we computed PPC between each of the electrodes in the left, right or back group as “seed” and all other electrodes. Subsequently, we averaged the PPC maps for all electrodes within a group to generate an average PPC map for each group. Figure 1d shows the average PPC maps with seed electrodes on the left (top) and right (bottom), which we aim to contrast across different age groups and disease conditions.

**Fig. 1:**
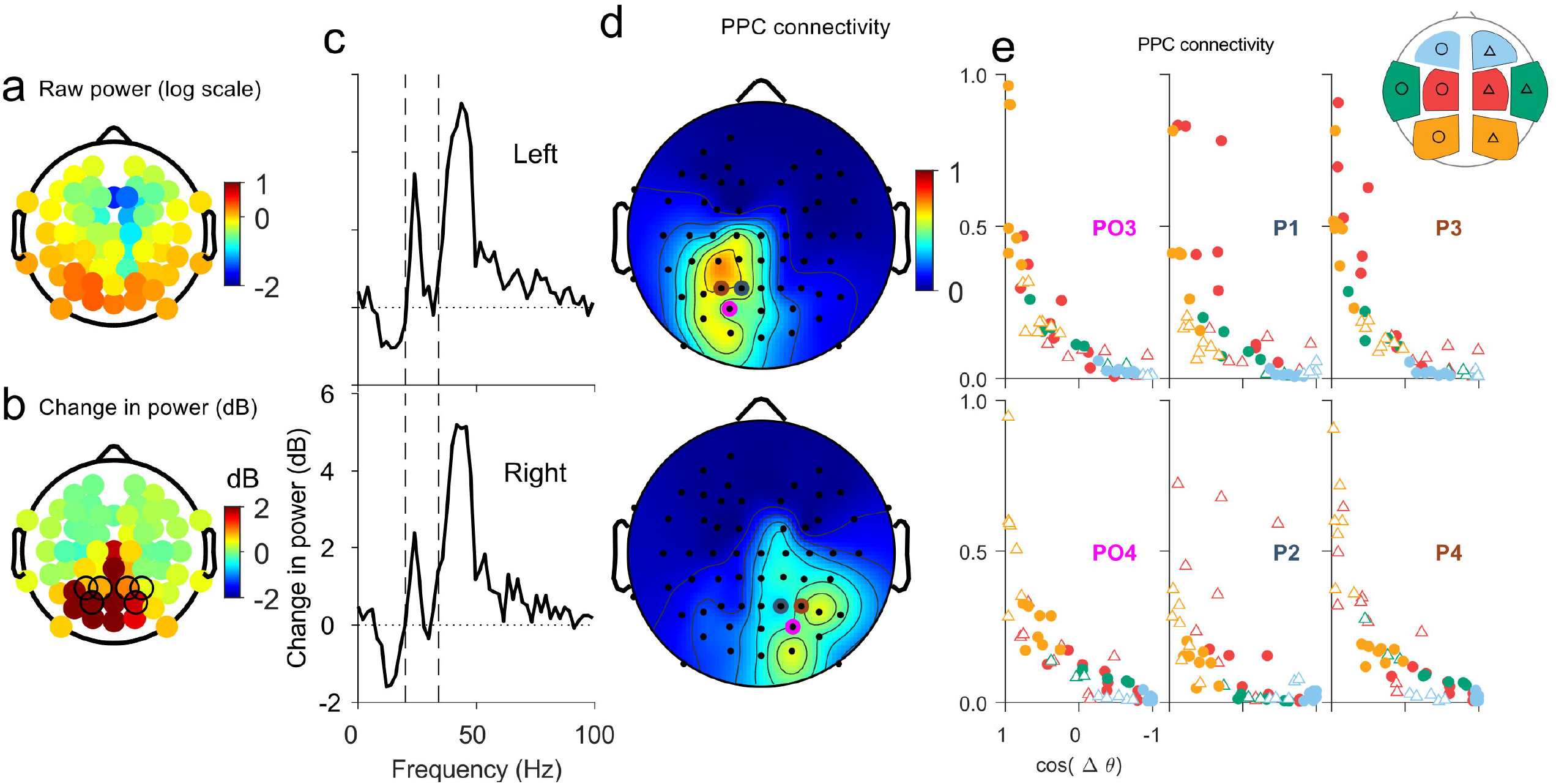
Power and Connectivity for a sample subject in slow gamma band across left and right seed electrode groups. (a) Average scalp map of absolute slow gamma band power (in log10 (µV^2^) units) during stimulus duration [0.25 0.75] s across 64 electrodes, for a representative subject. (b) Similar to (a) for change in slow gamma power computed relative to the baseline power, corresponding to the interval [-0.5 0] s, in decibels (dB). Note that the predominant power is in occipital electrodes in (a) and (b) scalp maps. (c) Change in power spectral density (PSD) across the left and right seed electrode groups. These electrode groups are shown in black circles in b. Dashed lines indicate the slow gamma range. (d) The scalp map of average FC measured using PPC between all the electrodes within each of the seed electrodes in the left (electrodes P3, P1, and PO3; top plot) and right (P2, P4, and PO4; bottom plot) groups. (e) Functional connectivity as a function of inter-electrode distance from the seed electrodes plotted for each of the three electrodes in the group with the other remaining electrodes. Schematic on top right shows the four electrode clusters (frontal, central, occipital, and temporal) defined over both the hemispheres, denoted by the four colours, cyan, red, yellow, and green respectively, with the hemisphere side denoted by filled circle for left and open triangle for right.

The maps in Figure 1d show that FC falls with increasing inter-electrode distance, as expected. However, some electrodes may have higher FC with respect to the seed electrodes than what is expected based on their inter-electrode distance. For example, left and right group of seed electrodes may show enhanced FC due to potential inter-hemispheric synchrony (Engel et al., 1991). Figure 1e shows the FC for different electrodes (frontal, central, temporal and occipital; see inset for details) as a function of the inter-electrode distance (see Methods for details) from each one of the seed electrodes in the left (top) or right (bottom) electrode group. The FC decrease with inter-electrode distance did not show any obvious deviations based on the brain region. In particular, the homologous inter-hemispheric occipital electrodes (yellow triangles and circles for top and bottom plots, respectively) did not show greater FC as compared to intra-hemispheric electrodes in the central region (red circles and triangles for top and bottom plots, respectively). Further, different seed electrodes within a group (different columns in Figure 1e) showed similar trends. We therefore considered FC values between electrodes pairs solely based on inter-electrode distance, ignoring which electrode group they belonged to (i.e., ignoring the color and symbol in Figure 1e), and further pooled across different seed electrodes (pooling data across the three columns in Figure 1e).

### Alpha and slow gamma FC shrinks with healthy aging

We first studied the dependency of alpha, slow gamma, and fast gamma FC on age by contrasting between the middle-aged (N=91) and elderly (N=127) groups. Figure 2a shows the median change in power (in the stimulus period from baseline) in alpha (top row), slow-gamma (middle) and fast-gamma (bottom row) for the two age groups. As shown previously (Murty et al., 2020), power in the alpha range remained unchanged whereas the power in slow gamma and fast gamma reduced across the age groups, primarily in the occipital electrodes. Figure 2b depicts the average FC maps for the middle-aged and elderly age groups over both the left and right occipital group of electrodes as seed for FC computation (FC map with right group is flipped horizontally since the individual maps for left and right electrode groups showed similar results; refer ‘visualizing average connectivity’ section in Methods for more details). This plot shows a subtle (but highly significant; as shown below) reduction in the FC of the elderly group compared to the middle-aged in the alpha range, which can be observed by comparing the red coloured contours that represent FC of 0.25. The shrinkage of this contour for the elderly group can also be observed in the gamma bands, although the effect is weaker.

**Fig. 2:**
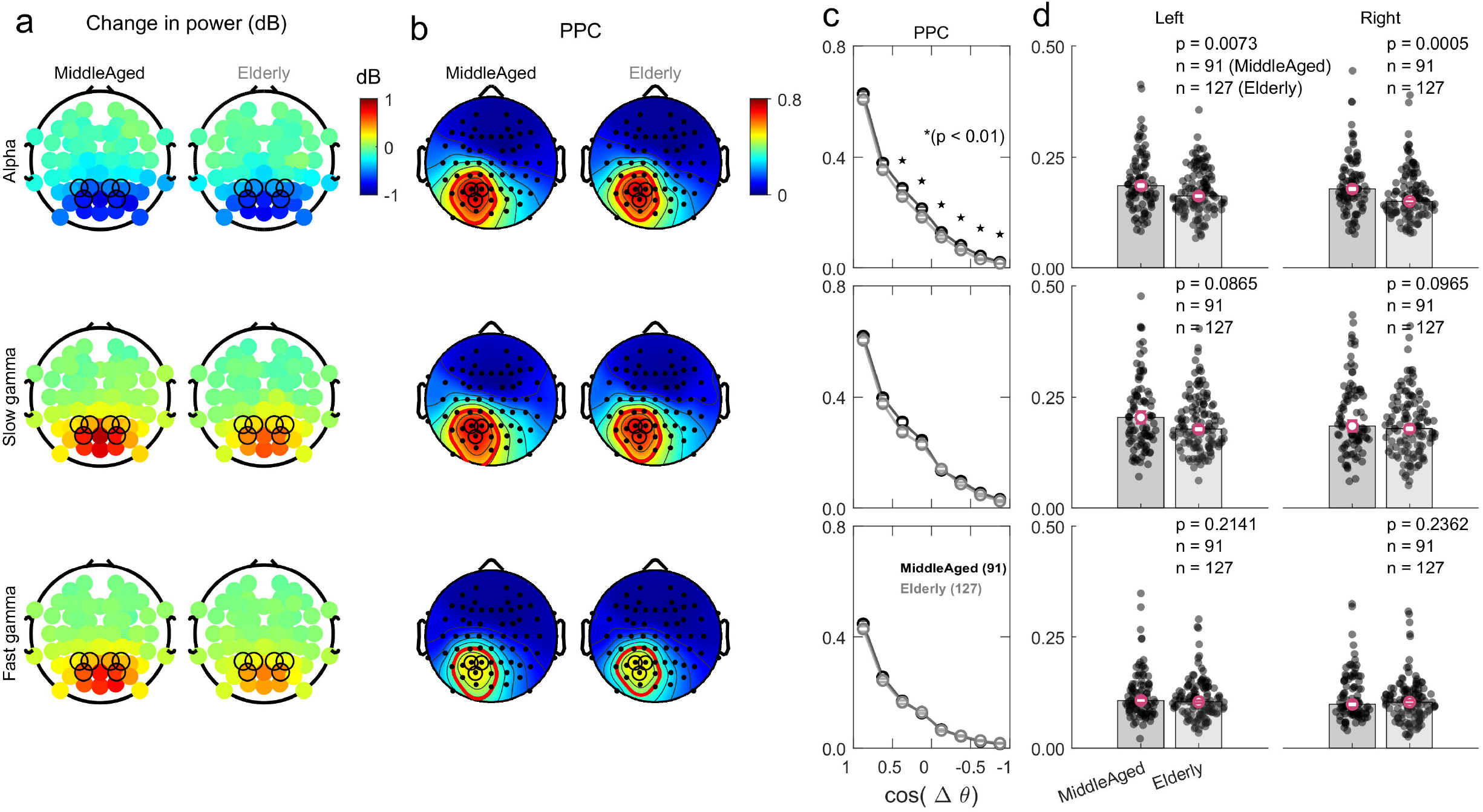
Effect of aging on FC. (a) Median change in power scalp plots in three frequency bands: alpha (top row), slow gamma (middle row), and fast gamma (bottom row) across middle-aged (N=91) and elderly (N=127) subjects. (b) Median PPC FC scalp map for the two age groups, averaged across the maps with the seed electrodes in the left and right group (after flipping around the vertical axis), along with 25% normalized FC contour in red. (c) PPC with inter-electrode distance plotted over linearly spaced bins in cosine of angular separation, averaged for each of the seed electrodes within left and right groups and then taking the median over subjects. Bins that are significantly different across middle-aged and elderly groups (p<0.01; KW test) are marked with an asterisk (*). (d) FC values averaged across bins with *cos*(Δ*θ*) in the range [-0.5 □ 0.5] for the two age-groups, separately for the right and left seed electrode groups. The p-values obtained using KW test across the age groups indicate prominent difference in the alpha band.

To quantify these differences, FC values for all electrode pairs falling within a specific inter-electrode distance range were averaged to yield a single FC versus distance plot for each electrode groups in each subject. Figure 2c shows the median FC versus distance plot for subjects within each age group, averaged over the left and right seed electrodes for each subject. FC for elderly subjects was significantly lower than middle-aged subjects in most of the bins in the alpha band, as indicated by the asterisks (p-value<0.01) in Figure 2c, top row. The FC values for elderly were lower than middle-aged for some bins in the slow-gamma band as well (middle panel), although the difference did not reach significance. For fast-gamma band (bottom plot), the differences were negligible.

We further averaged all FC values of electrode pairs within inter-electrode distance range of [-0.5 to 0.5] to yield a single FC value per electrode group per subject (Figure 2d; see Methods for details). This range was chosen to limit the dominance of high FC values close to seed electrode in the average which could be more affected by volume conduction. Figure 2d shows median FC values within the two age groups, separately for each seed electrode group. Both electrode groups showed significantly reduced alpha FC (top row) in the elderly over the middle-aged subjects (left group: *χ*^2^(1) = 7.21, n = 91/127, p = 0.007; Kruskal Wallis (KW) test; right group: *χ*^2^(1) = 11.98, n = 91/127, p = 0.0005; KW test). Similar findings were observed for the back electrode group as well (*χ*^2^(1) = 4.78, n = 91/127, p = 0.028; KW test; Figure 2 Supp Fig. 1). FC was not significantly different across the age groups in both the slow (middle row) and fast gamma (bottom row) bands (statistics are shown in the plots), although slow gamma FC almost reached significance (Figure 2d, second row).

The reduction in alpha FC was small in absolute terms (0.185 for middle-aged and 0.167 for elderly when all sides were pooled, or a reduction of 9.6%), potentially due to the method of averaging that was done over a wide range of inter-electrode distance with varying FC values. Nevertheless, the reduction was consistent across individual inter-electrode bins and electrode groups and therefore highly significant. We quantified the robustness of these results by estimating the Bayes factor (BF), which is the ratio of the marginal likelihood of the alternate hypothesis (higher FC in middle-aged than the elderly) and the null hypotheses (comparable FC in middle-aged and elderly), given the observed data (see Methods for details). BF values above 10 provide “strong” evidence in favor of the alternative, while BF above 3 provide “substantial” evidence (Jeffreys, 1998). For alpha FC, BF values were 20.42, 37.83 and 6.78 for the Left, Right and Back electrode groups, and 21.32 when electrode groups were pooled. For slow-gamma, BF values were 2.73, 2.31, 0.93 and 1.89 for left, right, back and combined groups, suggesting “anecdotal” evidence (see Methods for details). For fast gamma, these values were 0.90, 0.77, 0.33 and 0.67, suggesting insufficient evidence in favor of the alternate hypothesis.

To test whether FC results could be due to differences in power in the two groups, we performed a power-matching analysis. We first uniformly binned subjects from both middle-aged and elderly groups into several sub-groups based on their power, and subsequently randomly selected an equal number of subjects from each sub-group (see Methods for more details). This procedure ensured that the power distribution across the two age groups matched (thereby matching the average, median, and other statistics). Figure 3a shows the mean change in PSD for the left group of electrodes after performing the power matching procedure in alpha (top row, N=88 subjects in both age groups), slow-gamma (middle row, N=84) and fast-gamma bands (bottom row, N=84). The frequency bands based on which power matching was done is highlighted by dashed lines. Figure 3b and 3c shows the corresponding PPC maps and PPC versus distance plots for the left electrode group (note that unlike Figure 2b-c, left and right electrode groups cannot be averaged since different subjects are chosen for each side). Corresponding results for the back electrode group is shown in Figure 3 Fig. Supp 1.

**Fig. 3:**
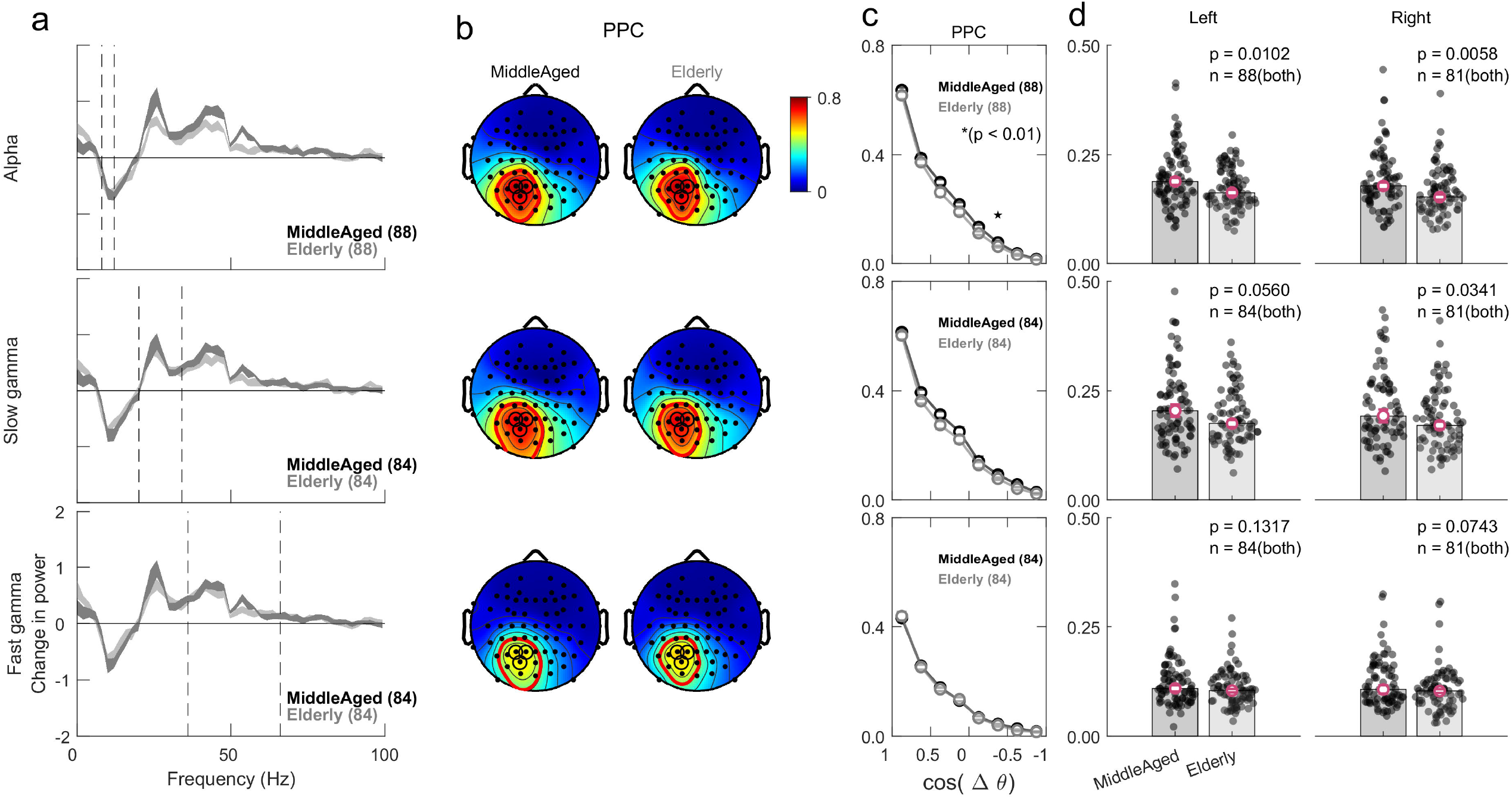
Effect of aging on FC after power-matching. (a) Change in PSDs across the middle-aged and elderly age groups after power matching applied over the left group of electrodes. Power matching was done separately in the particular frequency range (across the rows, from top: alpha, slow gamma, and fast gamma), as indicated by the dashed lines. The width of the PSD trace represents SEM across the subjects within the associated frequency bin. Note that different number of subjects were obtained for each frequency range (N=88, 84 and 84 for alpha, slow gamma and fast gamma, respectively). (b-c) Similar to Fig. 2b-c, but computed only for the selected subjects for the left electrode group (note that in this case left and right groups cannot be averaged since there are different subjects in each group). d) Same as 2d, but for sub-selected population for each electrode group. Note that the number of samples in both age groups match as a consequence of power matching.

Overall, the results remained similar after power matching. Alpha FC reduced with age (Left: *χ*^2^(1) = 6.6, n = 88, p = 0.01, BF = 17.33; Right: *χ*^2^(1) = 7.60, n = 81, p = 0.005, BF = 11.16; Back: *χ*^2^(1) = 3.09, n = 87, p = 0.07, BF = 2.44). Reduction in slow gamma FC also approached significance (Left: *χ*^2^(1) = 3.65, n = 84, p = 0.05, BF = 3.91, KW test, Right: *χ*^2^(1) = 4.49, n = 81, p = 0.03, BF = 4.45; Back: *χ*^2^(1) = 2.17, n = 78, p = 0.141, BF = 1.05), while fast gamma FC remained indifferent across age groups (Left: *χ*^2^(1) = 2.27, n = 84, p = 0.13, BF = 1.72, Right: *χ*^2^(1) = 3.18, n = 81, p = 0.07, BF = 1.94; Back: *χ*^2^(1) = 0.54, n = 79, p = 0.45, BF = 0.36). Similar results were obtained when we used coherence or PLV instead of PPC (data not shown).

Results after power matching varied slightly across different iterations since different subset of subjects could be selected within each age group, under the power distribution matching criteria. Figure 3 shows results of an iteration for which the p-values in Figure 3d were close to the median of the p-values computed over 50 iterations. The p-values for alpha, slow-gamma and fast-gamma FC were less than 0.05 for ∼96%, ∼56% and ∼18% of the iterations, implying consistent, borderline (weak) and insignificant reductions in FC in these three bands, respectively.

### Slow gamma connectivity spread shrinks with cognitive impairment, not for alpha

The MCI subjects (referred hereafter as cases) were compared with age (±1 year) and gender matched healthy subjects (referred to as controls) to study the effect of cognitive disorder on FC. As in our previous study (Murty et al., 2021), since the number of controls far exceeded the cases (see Methods for details), we averaged the relevant metrics (power, FC) across all age and gender matched controls for each case subject, culminating in the same number of cases and controls. Figure 4 and Figure 4 Supp 1 show the results in the same format as Figure 2 and Figure 2 Supp 1. Figure 4a illustrates notable reduction in occipital slow gamma and fast gamma power but not alpha power in cases over the respective control group, as reported previously (Murty et al., 2021). Although these power trends were similar to the effect of aging (Figure 2a), the corresponding FC results were different. Now, the highest reduction in FC was observed in the slow-gamma band, with negligible effect on alpha or fast-gamma bands. Reduction in slow gamma FC for the cases over controls was observed in all electrode groups, with BF values in the ‘anecdotal’ to ‘substantial’ range in spite of the small sample size (Left: w = 56, n = 11, *P*(*W* ≥ *w*) = 0.02, BF = 3.6, Wilcoxon signed rank (WSR) test); Right: w = 50, n = 11, *P*(*W* ≥ *w*) = 0.07, BF = 2.21, WSR test; Back: w = 58, n = 11, *P*(*W* ≥ *w*) = 0.01, BF = 10.67, WSR test). When all electrodes were averaged, the reduction was highly significant: (w = 59, n = 11, *P*(*W* ≥ *w*) = 0.009, BF = 6.87, WSR test). In contrast, alpha and fast-gamma FC values were not significantly different (statistics shown in the Figure), with BF rarely exceeding 1 (not shown).

**Fig. 4:**
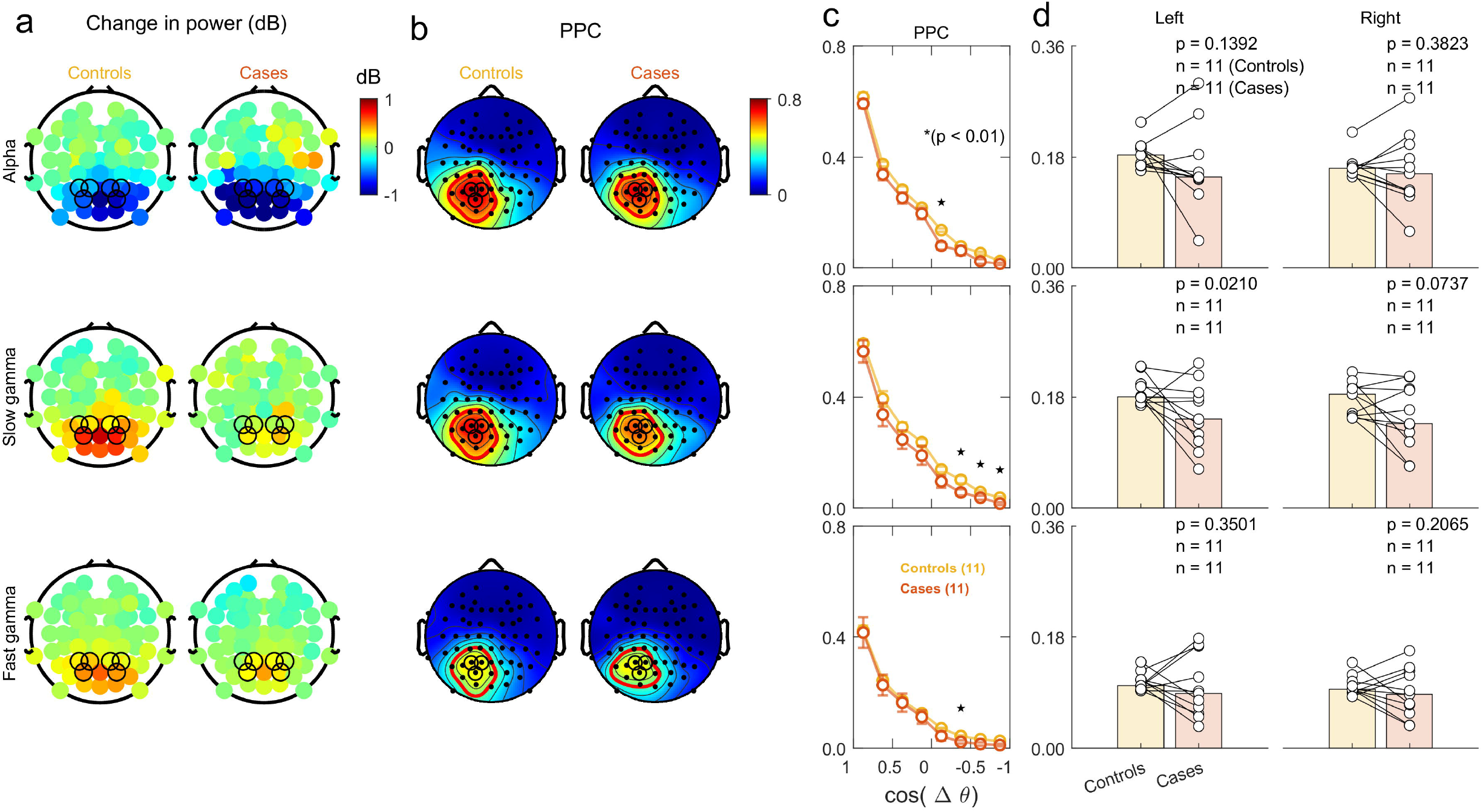
Effect of MCI on FC. (a) Change in power scalp plots for each frequency band over rows, averaged across MCI (case) subjects (N=11) and corresponding age and gender matched control subjects. Although there were many control subjects for each case, their relevant metrics were averaged to yield a single control data point for each case. (b-c) Scalp maps and connectivity profiles for cases and controls; similar format as Fig 2b-c. (d) Bar plots indicate the median FC over the defined inter-electrode distance range (similar to Fig. 2d) for the cases and controls. Individual case and control points are shown as connected white circles.

Figure 5 shows the same results after choosing a single control subject for each case subject with minimal difference in band power (note that different control subject may get chosen for the same case subject in different frequency bands and seed electrode group). Interestingly, the reduction in slow-gamma FC was more prominent now, with BF values in the “strong” range (Left: w = 59, n = 11, *P*(*W* ≥ *w*) = 0.009, BF = 10.73, WSR test; Right: *w* = 56, n = 11, *P*(*W* ≥ *w*) = 0.02, BF = 3.71, WSR test; Back: *w* = 66, n = 11, *P*(*W* ≥ *w*) = 0.0005, BF =17.65, WSR test; Overall: *w* = 60, n = 11, *P*(*W* ≥ *w*) = 0.007, BF = 10.47, WSR test), while no differences were observed in alpha and fast-gamma FC.

**Fig. 5:**
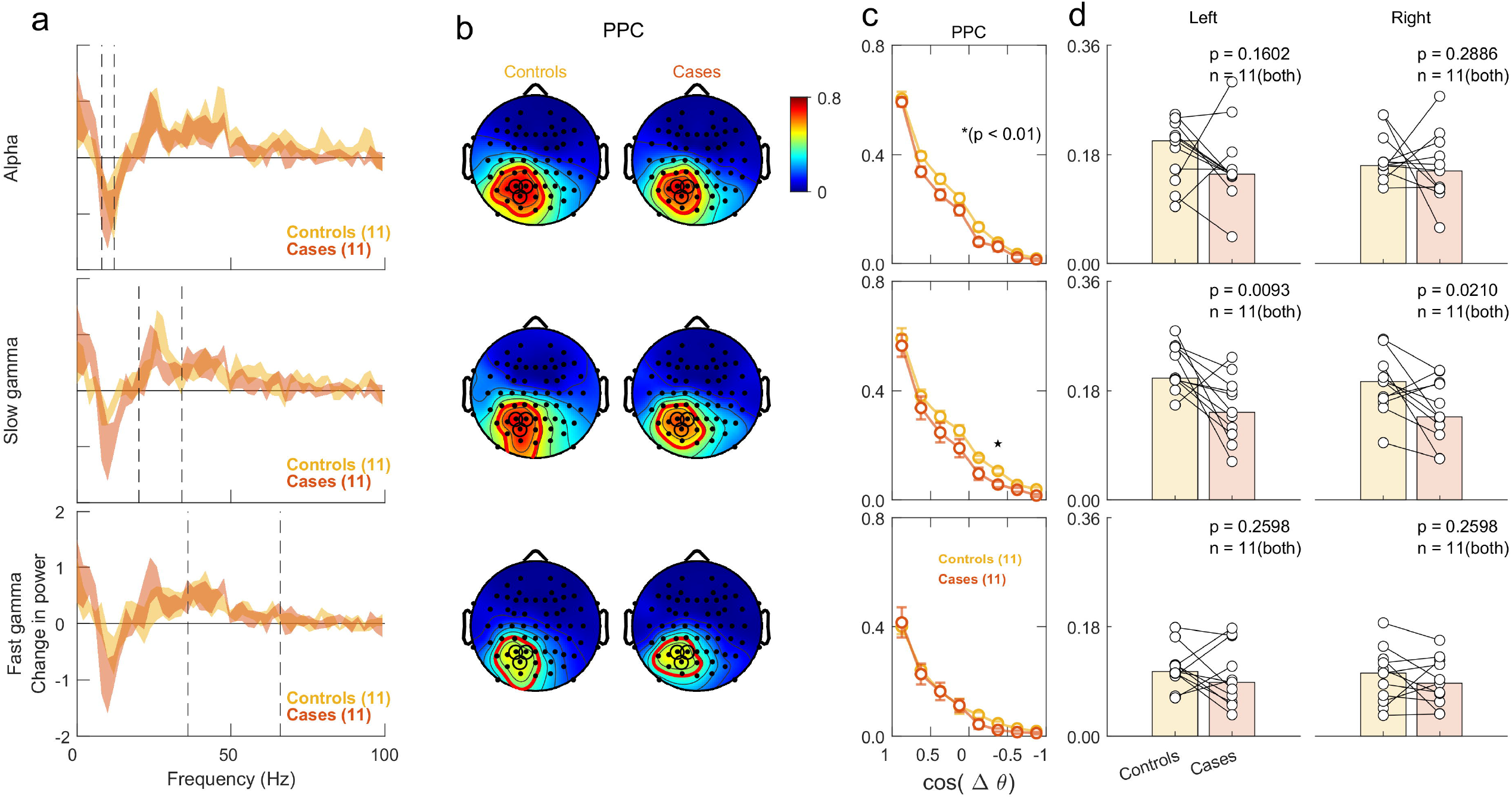
Effect of MCI on FC after power matching. (a) Change in PSDs averaged across case and controls subjects after power matching, which is done by taking one control subject for each case subject whose power is closest to the case subject. Note that this control subject may be different for different frequency ranges and seed electrode groups. (b-c) Scalp maps and connectivity profiles for case subjects and power-matched controls; similar format as Fig. 2b-c. Note that here data from left and right electrode groups could be averaged since both had the same number of subjects. (d) Same as Fig. 4d.

### Dependence of FC on power and age

The dependence of FC on age and change in power was further explored with a robust linear regression model applied over all the healthy subjects (refer Methods section) (Figure 6). Alpha, slow gamma, and fast gamma FC was individually regressed with subject age and the change in band power as independent variables, separately for each seed electrode group. Figure 6a shows the scatter plot of alpha, slow gamma, and fast gamma FC versus subject age, while Figure 6b shows the same versus change in power, for the left electrode group (similar results were obtained for other groups; data not shown). The coefficient in the fitted model and its significance (p-value) are also shown in the plots. The p-value relates to the t-statistic of the two-sided hypothesis test (*H: b* = 0), given the other terms in the model. Consistent with previous results, alpha FC depended only on the age factor (*b*_*age*_ = -0.002; p-value = 0.002) and not on the change in power (*b*_*power*_ =0.004; p-value = 0.380). This can also be observed in the median alpha FC values binned in 5-year intervals as shown in black traces in Figure 6a, which decreased with age (left plot).

**Fig. 6.**
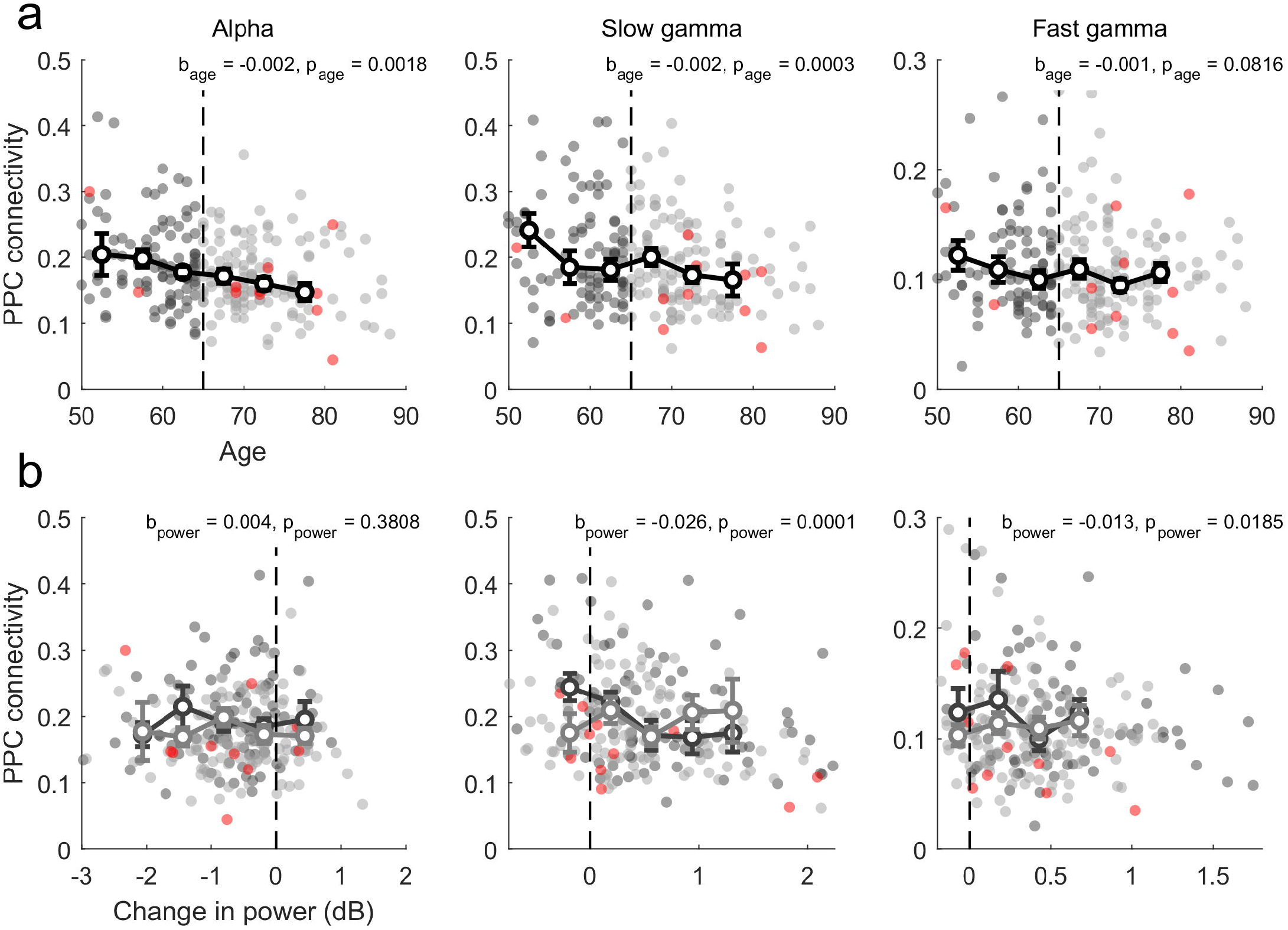
Dependence of FC on age and power. (a) FC against age for all subjects, coloured according to the subject category, namely middle-aged (dark grey), elderly (light grey) and MCI (red). The dependency of FC on age was considered in various frequency band i.e., alpha, slow gamma, and fast gamma, plotted across columns. The vertical dashed line marks the age 65, that separates middle-aged and elderly age groups. The representative errorbar depicts median FC, within uniformly spaced bins of 5 years width. The errorbar conveys the bootstrapped median SEM. (b) FC against the change in power in the respective frequency ranges, arranged similar to (a). The median FC errorbars for each bin, with 0.6 dB width, were plotted for middle-aged and elderly groups separately. The errorbar conveys the bootstrapped median SEM. The linear regression coefficients, with the respective p-value of significance are indicated for age and power variables for each of the frequency bands. Only healthy subjects were considered for regression analysis.

In contrast, regression analysis for slow gamma revealed additional dependencies not observed earlier. In particular, slow gamma FC was found to depend on both, age (*b*_*age*_ = - 0.002; p-value = 0.0003) and change in power (*b*_*power*_ =-0.026; p-value = 0.0001). Unexpectedly, FC was negatively correlated with change in power (potential reasons are discussed below). The dependence of slow-gamma FC on age can also be observed in the median FC values binned over 5-year intervals (black trace in Figure 6a). This dependence was weaker in our previous analysis because we had partitioned the subjects into the two age groups, middle-aged and elderly. Indeed, when we performed the same regression analysis but after replacing the age with age category (0 for middle aged and 1 for elderly) and excluding the power dependence, the resulting coefficient barely reached significance in the slow gamma band (*b*_*age*_ = -0.022; p-value = 0.03) even though it remained significant for the alpha band (*b*_*age*_ = -0.024; p-value = 0.0028). Results for fast gamma FC were largely consistent with previous analyses, showing no dependence on the age factor (*b*_*age*_ = -0.001; p-value = 0.0816). In addition, there was a weak dependence on the change in power (*b*_*power*_ =-0.013; p-value = 0.0185), similar to the dependence observed for slow gamma.

The MCI subjects shown in red showed the expected results: these points were scattered across either side of the black traces for alpha and fast-gamma (left and right plots in Figure 6a), but most of the points lay below the black trace for slow-gamma (middle plot).

## Discussion

We studied the change in stimulus-induced EEG FC, computed using PPC, with aging and cognitive impairment in elderly (>49 years) human subjects. With healthy aging, although power reduced in slow and fast gamma bands but not alpha band (Murty et al., 2020), FC reduced in alpha and slow-gamma but not in fast-gamma band. These results remained consistent even when power in the two age groups were matched or accounted for using multiple linear regression. On the other hand, cognitive decline due to MCI reduced power in both slow and fast gamma but not alpha band (Murty et al., 2021), while FC reduced only in the slow gamma band. The reduction in slow gamma FC was observed even when power was matched for the two age groups. These results remained consistent across the left, right, and back electrode groups and over other FC measures, such as coherence and PLV. This indicates a potential role of induced EEG FC in detecting early onset of MCI.

### Relationship between FC and power

We found a negative correlation between PPC connectivity and change in power in slow and fast gamma bands (Figure 6b). The reasons are unclear, but in line with other studies that have reported similarities between PSDs and network measures computed from FC measures such as PLV, and cautioned against treating them as completely separate measures (Demuru et al., 2020). One possibility is that when the oscillatory signal is weak, the FC is governed by the background spectral power which may have a larger spatial spread. Another possibility is that the source of the oscillation is deeper, which would lead to lower power at the sensor level but nonetheless stronger FC across sensors since more sensors would capture the source activity.

On the other hand, several reports have shown that power and FC capture different features of the underlying neural activity. For example, differences in power versus FC with age and cognitive decline (MCI) reported here have also been observed in resting state EEG, although the strongest effect is observed at lower frequencies such as delta and theta bands (Meghdadi et al., 2021). Similarly, FC could convey different information compared to PSDs about mental disorders. For example, patients with schizophrenia have higher EEG complexity at lower frequencies (Takahashi et al., 2010), which is shown to reflect reduced FC (Friston, 1996). In addition, FC is shown to capture subject specific information like the demographic variables, while being consistent across a variety of tasks (Nentwich et al., 2020). However, a recent study reported source and sensor space FC to be less reproducible than either the absolute or relative powers (Duan et al., 2021). Note that most of these earlier studies used resting state EEG data. Our study extends these findings for stimulus-induced oscillations and shows that power and FC alter in distinct ways with aging and cognitive disorders, and the changes in FC can be observed even after accounting for the difference in power. Indeed, some studies have shown that compared to resting-state EEG, a paradigm involving visual stimulation/perception is more sensitive to AD disease effects (Barzegaran et al., 2016).

### Previous studies on the effect of aging and AD on power and FC

Alpha FC has been shown to be more reproducible than other frequencies (Duan et al., 2021), allowing high subject identifiability in resting state MEG, especially in the visual subnetwork (Sareen et al., 2021). Consequently, the reduction in alpha FC with aging has been demonstrated using different recording setups (MEG, EEG) and techniques. For example, a recent MEG study showed that FC in Upper Alpha (UA) band (10 – 14 Hz) falls with age (Pathak et al., 2022), owing to the reduction in individual peak alpha frequency (IAPF) with age (18 – 88 yrs), even though the FC (PLV and Phase lag index (PLI)) at the peak alpha frequency remained invariant. This matches with the falling trend of alpha (8 – 12 Hz) FC with age in our data (Figure 6a). Another study in eyes-closed resting state EEG (Scally et al., 2018) reported reduced alpha power and FC in the conventional UA band (10 – 12 Hz) in older adults compared to young adults, although power and FC at IAPF were similar in the two age groups. Note that our results are not inconsistent with previous studies that have shown reduction in alpha power (Scally et al., 2018) and IAPF with aging (Pathak et al., 2022), since these studies used a much wider age distribution than ours. We have previously shown that alpha power was indeed significantly less in middle-aged subjects (50-64 years) compared to young subjects between 20-49 years (Murty et al., 2020). Further, since we computed the PSDs over 500 ms of data, the resulting frequency resolution of 2 Hz was not enough to study changes in IAPF. Another study in resting state EEG (Moezzi et al., 2019) showed reduction in alpha FC (measured using imCOH) within old adults compared to the young, along with gamma FC, especially within the connections involving the occipital electrodes.

With AD progression, Dai and colleagues showed prominent disruption in FC in long-range connections (Dai et al., 2015). These long range connections across the brain have been proposed to play an important role in aiding human cognitive function (Bullmore and Sporns, 2012). In a MEG study, Berendse and colleagues showed reduced fronto-parietal FC in 2 to 22 Hz range in AD patients over controls (Berendse et al., 2000). FC measures like phase coherence and amplitude correlation have been shown to provide insights into large-scale neuronal interactions, which degrade in neuropsychiatric disorders (Siegel et al., 2012).

### Neural mechanisms underlying changes in power and FC

Reduction in alpha FC has been attributed to various changes in brain morphology, like white matter hyperintensities (Quandt et al., 2020), cerebral reorganization or dedifferentiation processes (Edde et al., 2021). Recently, Ranasinghe and colleagues measured the PSDs of AD subjects and healthy controls and found enhanced delta/theta but reduced alpha/beta power, which they fitted using a neural mass model by appropriately changing the excitatory and inhibitory time-constants (Ranasinghe et al., 2022). They also measured regional tau and Aβ in AD patients by positron emission tomography. Surprisingly, excitatory time-constants were tightly correlated with tau levels, while inhibitory time-constants tightly correlated with Aβ depositions.

The excitatory-inhibitory interactions can also produce gamma oscillations (Wang, 2010). While fast gamma oscillations are thought to be due to parvalbumin-positive GABAergic interneurons (Bartos et al., 2007), slow gamma could be related to long-range inhibitory somatostatin interneurons, as reported in a mouse V1 study (Veit et al., 2017). The GABAergic system gets altered with AD pathogenesis, giving rise to E/I imbalance (Petrache et al., 2019). Indeed, cognitive functions in AD patients and mouse models can be improved by the correction of E/I imbalance though drug intervention (Bi et al., 2020). Dysfunction of these inter-neuronal networks with mental disorders could lead to lower gamma power and FC across areas (Uhlhaas and Singer, 2006).

The mechanisms discussed above are likely to reduce both power and FC in different frequency bands. How can then a reduction in alpha FC without changes in power or vice versa be explained? One possibility could be due to the presence of a variety of neuro-compensatory mechanisms that may cause power and FC to change differently. For example, Pathak and colleagues (2022) showed that the effect of increased axonal transmission delays due to degeneration of white matter tracts with aging can be cancelled by enhancing inter-areal coupling (Pathak et al., 2022), leading to FC at IAPF that is invariant with age. Similarly, Hinkley and colleagues (2011) reported unchanged alpha power but reduced FC (measured through PLV) in Schizophrenia patients compared to controls, which they attributed to disruption in long range connections but intact local connections.

### Comparison of FC measures

FC measures can be broadly classified into methods that are sensitive to volume conduction (such as coherence, PLV and PPC) versus those that are not (for example, imaginary part of coherence (Nolte et al., 2004), phase lag index (PLI; (Stam et al., 2007))). We used PPC in this report since the measures that are insensitive to volume conduction also tend to be less reproducible (Duan et al., 2021). Since our main aim was to test whether stimulus-induced gamma FC could be modulated by aging or cognitive impairment, we used the measure that is widely used and has been shown to yield reproducible results (Duan and colleagues found that PLV had highest reproducibility, and PPC is an unbiased estimator of the square of PLV). Further, the potential influence of volume conduction was reduced by powr-matching, which discards the common power factor (van Vliet et al., 2018). In addition, to exclude the local connections that are corrupted by volume conduction, we considered average FC over the electrode pairs within [-0.5 to 0.5] in *cos*(Δ*θ*) space. Future studies that incorporate other FC measures that are insensitive to volume conduction or incorporate this analysis in the source space will be useful to reveal the underlying neural mechanisms behind the differences in power and FC with aging and cognitive impairment.

## Materials and Methods

### Human subjects

The EEG dataset was collected from 257 human subjects (females: 106) aged 50 to 88 years as part of the Tata Longitudinal study of aging (TLSA), of which usable data was obtained from 244 subjects (227 healthy, 12 MCI and 5 AD; see Murty et al., 2021 for detailed selection criteria). Subjects were recruited from urban Bengaluru communities and were evaluated by trained psychiatrists, psychologists and neurologists affiliated with National Institute of Mental Health and Neuroscience (NIMHANS) and M.S. Ramaiah Hospital, Bengaluru. Cognitive performance was evaluated using ACE (Addenbrooke’s Cognitive Examination), CDR (Clinical Dementia Rating) and HMSE (Hindi Mental State Examination) tests (see Murty et al., 2021 for more details). The AD group was very small, and most subjects did not have appreciable gamma, and were therefore not considered for further analysis. We further discarded 10 subjects (9 healthy and 1 MCI) because data was collected using 32 channels. All results are based on the remaining 218 healthy and 11 MCI subjects.

All subjects took part against monetary compensation and provided signed informed consent. All the procedures were approved by The Institute Human Ethics Committees of Indian Institute of Science, NIMHANS, Bengaluru and M.S. Ramaiah Hospital, Bengaluru.

### Experimental setting and behavioural task

Experimental setup and details of recordings have been explained in detail previously (Murty et al., 2020, 2021) and are briefly summarised here. EEG was recorded from 64-channel active electrodes (actiCap) using BrainAmp DC EEG acquisition system (Brain Products GMbH) and were placed according to the international 10-10 system. Raw signals were filtered online between 0.016Hz (first-order filter) and 1kHz (fifth order Butterworth filter), sampled at 2.5kHz, and digitized at 16-bit resolution (0.1µV/bit). It was subsequently decimated to 250 Hz. Average impedance of the final set of electrodes was (mean±SEM) 7.82 ± 0.02 kΩ. EEG signals recorded were in reference to electrode FCz during acquisition (unipolar reference scheme).

All subjects sat in a dark room facing a gamma-corrected LCD monitor (dimensions: 20.92 × 11.77 inches; resolution: 1289 × 720 pixels; refresh rate: 100 Hz; BenQ XL2411) with their head supported by a chin rest. It was placed ∼58 cm from the subjects and subtended 52° × 30° of visual angle for full screen gratings.

In the experiment, subjects performed a visual fixation task, wherein 2-3 full screen grating stimuli were presented for 800ms with an inter-stimulus interval of 700ms after a brief fixation of 1000ms in each trial using a customized software running on MAC OS. The stimuli were full contrast sinusoidal luminance achromatic gratings with one of the three spatial frequencies (1, 2, and 4 cycles per degree (cpd)) and four orientations (0°, 45°, 90°, and 135°).

### Artifact Rejection

We implemented a fully automated artifact rejection framework (for details, see (Murty et al., 2020, 2021; Murty and Ray, 2022), in which outliers were detected as repeats with deviation from the mean signal in either time or frequency domains by more than 6 standard deviations, and subsequently electrodes with too many (30%) outliers were discarded. Subsequently, stimulus repeats that were deemed bad in more than 10% of the other electrodes (see below) or in the visual electrodes were discarded. This gave rise to a set of good electrodes and common bad repeats for each subject. Overall, this led us to reject 15.33±0.36% (mean±SEM) stimulus repeats. We also rejected electrodes with high impedance (>25kΩ). Finally, as in our previous studies, we computed slopes for the baseline power spectrum between 56-84Hz range for each unipolar electrode and rejected electrodes whose slopes were less than 0. Based on these criteria, 15.30±0.72% (mean±SEM) electrodes were labelled bad. Some subjects who had predominantly bad electrodes (>40 electrodes) were rejected later in the pipeline.

Eye position was monitored using EyeLink 1000 (SR Research Ltd), sampled at 500Hz. Any eye-blinks or change in eye position outside a 5° fixation window during -0.5s to 0.75s from stimulus onset were noted as fixation breaks, which were removed offline. This led to a rejection of 16.54±0.059% (mean±SEM) stimulus repeats. Overall, we rejected (mean±SEM) 31.88±0.81% of the repeats.

### EEG data analysis

All analyses were performed using unipolar reference scheme. The 10 occipital electrodes considered for analysis were separated in the three groups: P3, P1, PO3 (left group); P2, P4, PO4 (right group) and POz, O1, Oz, O2 (back group). These electrodes were considered as seed electrodes in the connectivity computation.

All the data analyses were done using custom codes written in MATLAB (MathWorks. Inc; RRID:SCR_001622). Power spectrum and FC measures were obtained using the Fieldtrip Toolbox ((Oostenveld et al., 2010), RRID:SCR_004849). For spectral analysis, like our previous study (Murty et al., 2020), we chose [-500 0] msec as the baseline period and [250 750] msec as the stimulus period, with 2Hz resolution. FC was computed in the same stimulus period to avoid stimulus-onset transients. We chose PPC (Vinck et al., 2010) as a measure of FC since it is a consistent estimator and unbiased by the finite sample size bias, and independent of the distributions of relative phases (see Vinck et al., 2010 for more details). Results using coherence and PLV yielded similar results.

Change in power between stimulus period and baseline period for a frequency band was computed using the following formula

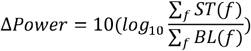

where ST is the stimulus power spectra and BL is the baseline power spectra, both averaged within relevant frequency bins (*f*), across all the analyzable repeats. Power was computed in three frequency bands: alpha (8-12 Hz), slow gamma (20-35 Hz), and fast gamma (36-66 Hz). These were averaged across all the three electrode groups. Scalp maps were generated using the *topoplot* function of EEGLAB toolbox ((Delorme and Makeig, 2004), RRID:SCR_007292).

### Visualizing average connectivity

Bilateral FC was computed between a pair of electrodes, which depended on the distance between the electrodes. Following previous literature (Kayser and Tenke, 2015), the inter-electrode distance was computed in angular space, as the Euclidean distance between the two locations on the scalp, in elevation and azimuth angle space. Specifically, we estimated the inter-electrode distance as *cos*(Δ*θ*), with Δ*θ* being the angular separation between the seed electrode and the electrode under consideration. We estimated the FC with inter-electrode distance profile from the seed electrode to the neighboring electrodes, by averaging FC between electrode pairs within a defined inter-electrode distance intervals (angular separation of [60□ □ 120□] or [-0.5 □ 0.5] in *cos*(Δ*θ*) space) covering the range of -1 to 1 with 0.25 bin width.

The FC was averaged across the left and right electrode groups for better visual representation in Figure 2b, 4b and 5b and the FC versus inter-electrode distance plots shown in 2c, 4c and 5c. The scalp maps were averaged by taking the mean of the FC map of the left electrode group and the mirror image of the FC map of the right electrode group. The FC versus distance profiles were simply averaged across the two electrode groups.

### Power-matching

Subjects were sub-selected from each age group, such that the distribution of change in band power within the groups were matched. We first binned the subjects within each age group with respect to the power and computed the greatest common distribution. Within each power bin, subjects were randomly selected from the group with more subjects, to match the subjects in the other group. Ultimately, the subjects in each bin were adjusted to arrive at the common distribution for the two age groups. This procedure eliminates the dependence of power on the effect under study, i.e., the FC (Churchland et al., 2010).

Since each iteration produced a different subset of subjects, the resultant p-value (obtained using Kruskal-Wallis Test) varied with each iteration. We executed this random sampling 50 times and chose a representative iteration (Figure 3) that represented the overall distribution of results (p-values). Specifically, we used the iteration with p-value near the median of the distribution for both left and right electrode groups.

Power matching for MCI vs. healthy comparison was done on a single subject basis. The subject among the age (within ±1 yr gap) and gender matched controls with minimal separation in change in power from the case subject was selected for each case.

### Linear Regression

The connectivity spread measure was modelled to linearly vary with age and change in power. This was done separately for each frequency band and seed electrode group. Robust linear regression was used to minimize the effect of outliers (Anon, 2009), using the ‘fitlm’ function in MATLAB.

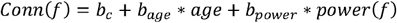

The p-value for each of the coefficients was computed for the t-statistic of the two-sided hypothesis test (*H: b* = 0), given the other terms in the regression model.

### Statistical Analysis

Kruskal-Wallis or Wilcoxon signed rank tests were used to compare medians over the two subject groups, and two-sample t-test or paired-sample t-test were used to compare means.

### Bayes factor analysis

Bayes Factor (BF) refers to the ratio of the marginal likelihood of alternate hypothesis to the likelihood of null hypothesis, given the data. The measure compares the alternate against the null hypothesis, rather than limiting the inference from the null hypothesis evidence (Jarosz and Wiley, 2014). In the present study, we used the Bayesian paired t-test with a Cauchy prior scaled to 1. Under this scheme, BF values below 1 suggests the absence of effect and evidence in favor of the null hypothesis, BF values between 1 and 3 provide anecdotal evidence in favor of the alternate hypothesis, while values between 3 and 10 provide substantial evidence and values above 10 provide strong evidence in favor of the alternative (Jeffreys, 1998). We calculated BF for right-tailed paired t-test (connectivity(controls) > connectivity(cases)) or (connectivity(Middle-aged) > connectivity(elderly)) for all the frequency bands. We used the MATLAB toolbox on BF by Bart Krekelberg (Krekelberg, 2021) based on (Rouder et al., 2012).

## Supplementary Figure legend

**Figure 2_Supplementary Figure 1**: Similar to Figure 2, with back electrode group as the seed for FC computation. (a-c) Change in power scalp map, FC maps and corresponding inter-electrode distance plots, similar to Fig. 2a-2c but only for the back electrode group. (d) The FC averaged over the predefined inter-electrode distance range, for the back group and all electrode groups condition, which was obtained by averaging the FC across all the 10 electrodes in the three seed electrode groups (left, right and back).

**Figure 3_Supplementary Figure 1**: Similar to Figure 3, with back electrode group as the seed for FC computation. Note that there is no ‘all’ condition here since the three electrode groups had different number of subjects.

**Figure 4_Supplementary Figure 1**: Similar to Figure 2_Supplementary Figure 1, for the case versus control comparison.

**Figure 5_Supplementary Figure 1**: Similar to Figure 5, with back electrode group as the seed for FC computation.

