## Supplementary Figures for "Healthy aging and cognitive impairment alter EEG functional connectivity in distinct frequency bands"

**Supplementary Figure legend**

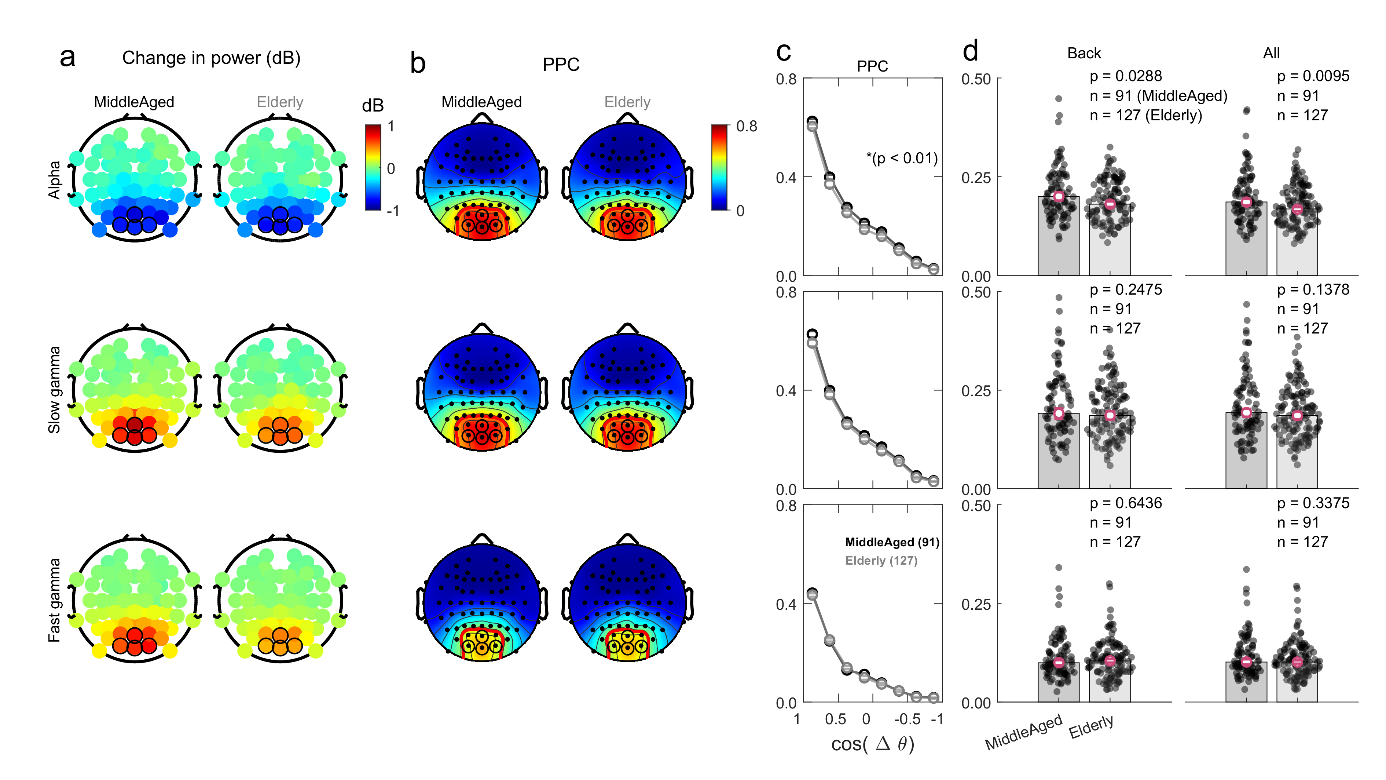

**Figure 2_Supplementary Figure 1**: Similar to Figure 2, with back electrode group as the seed for FC computation. (a-c) Change in power scalp map, FC maps and corresponding inter-electrode distance plots, similar to Fig. 2a-2c but only for the back electrode group. (d) The FC averaged over the predefined inter-electrode distance range, for the back group and all electrode groups condition, which was obtained by averaging the FC across all the 10 electrodes in the three seed electrode groups (left, right and back).

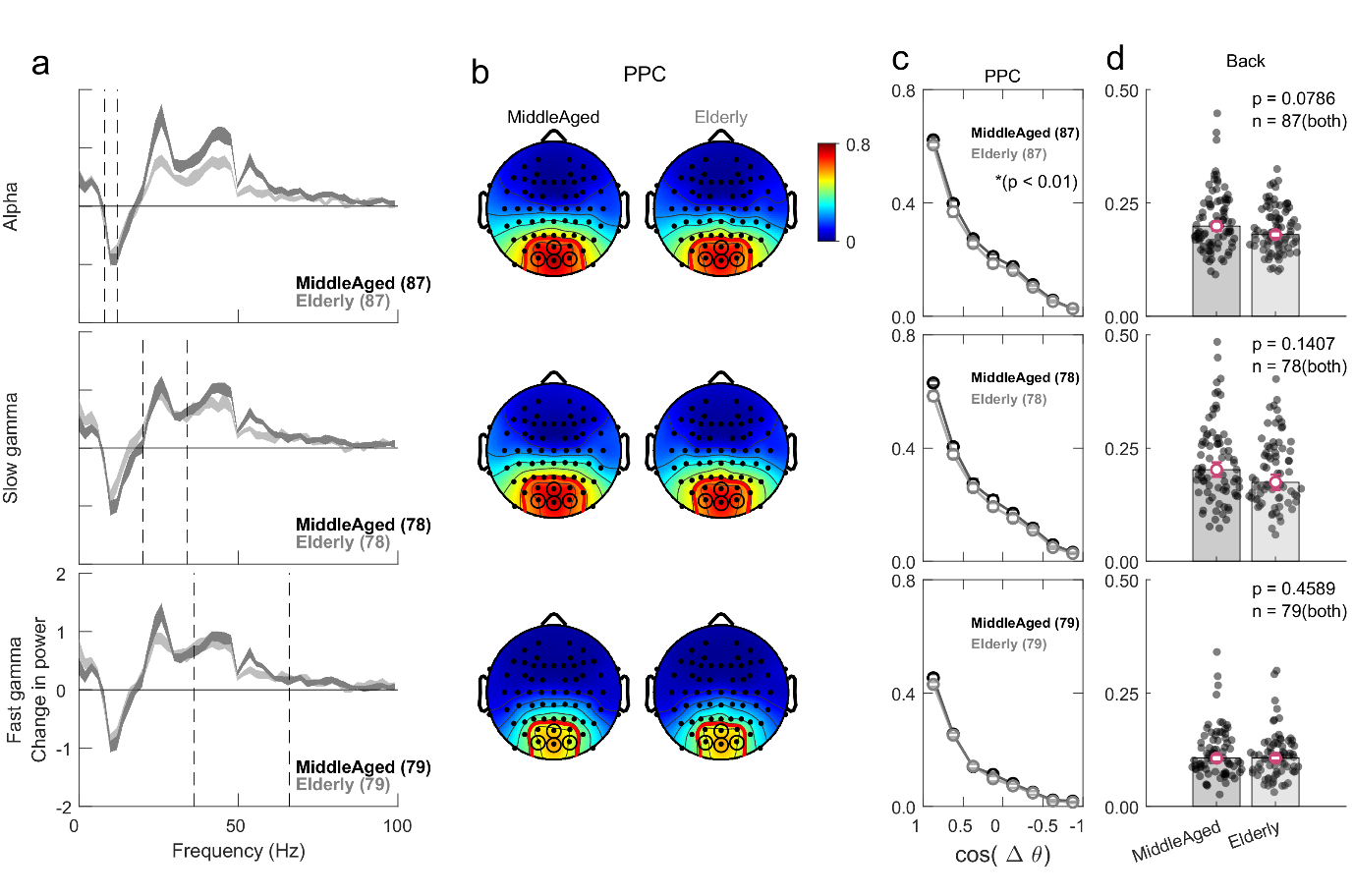

**Figure 3_Supplementary Figure 1**: Similar to Figure 3, with back electrode group as the seed for FC computation. Note that there is no ‘all’ condition here since the three electrode groups had different number of subjects.

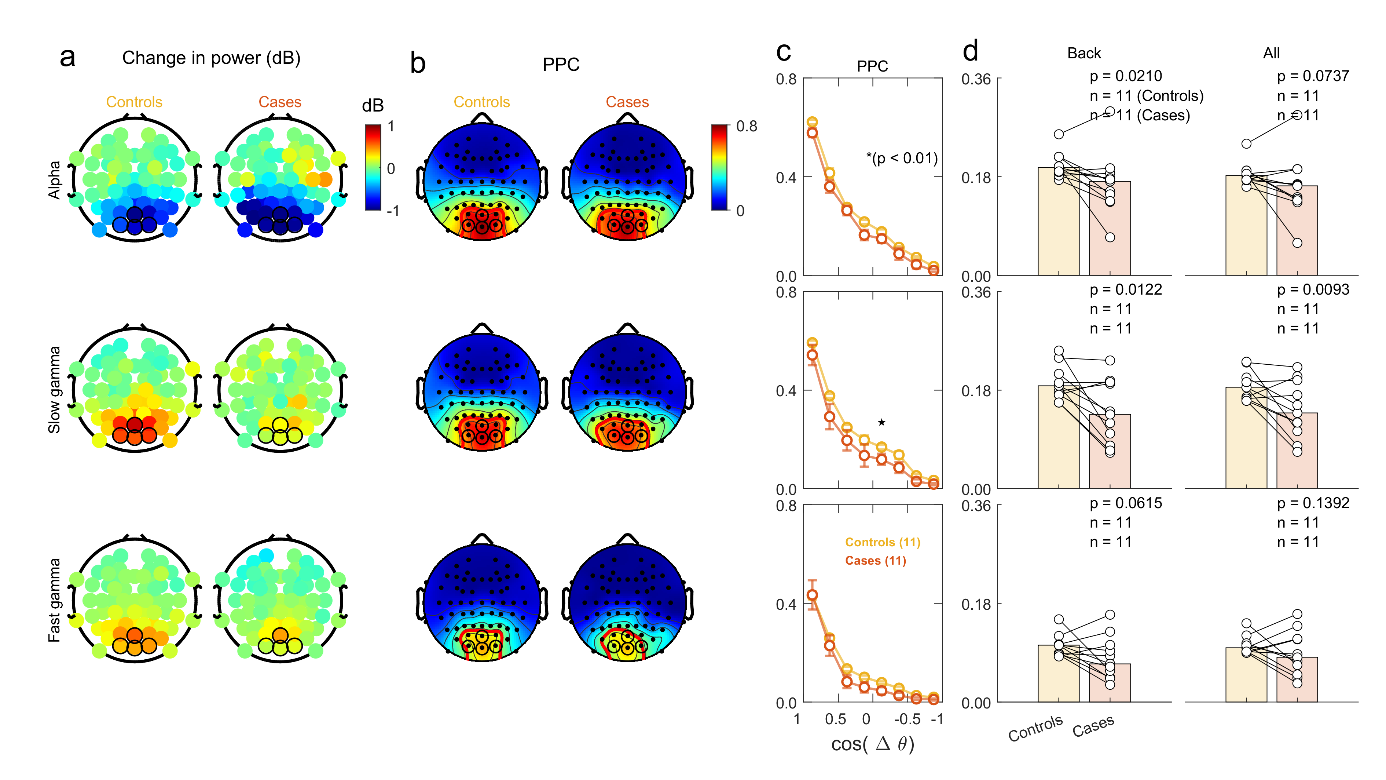

**Figure 4_Supplementary Figure 1**: Similar to Figure 2_Supplementary Figure 1, for the case versus control comparison.

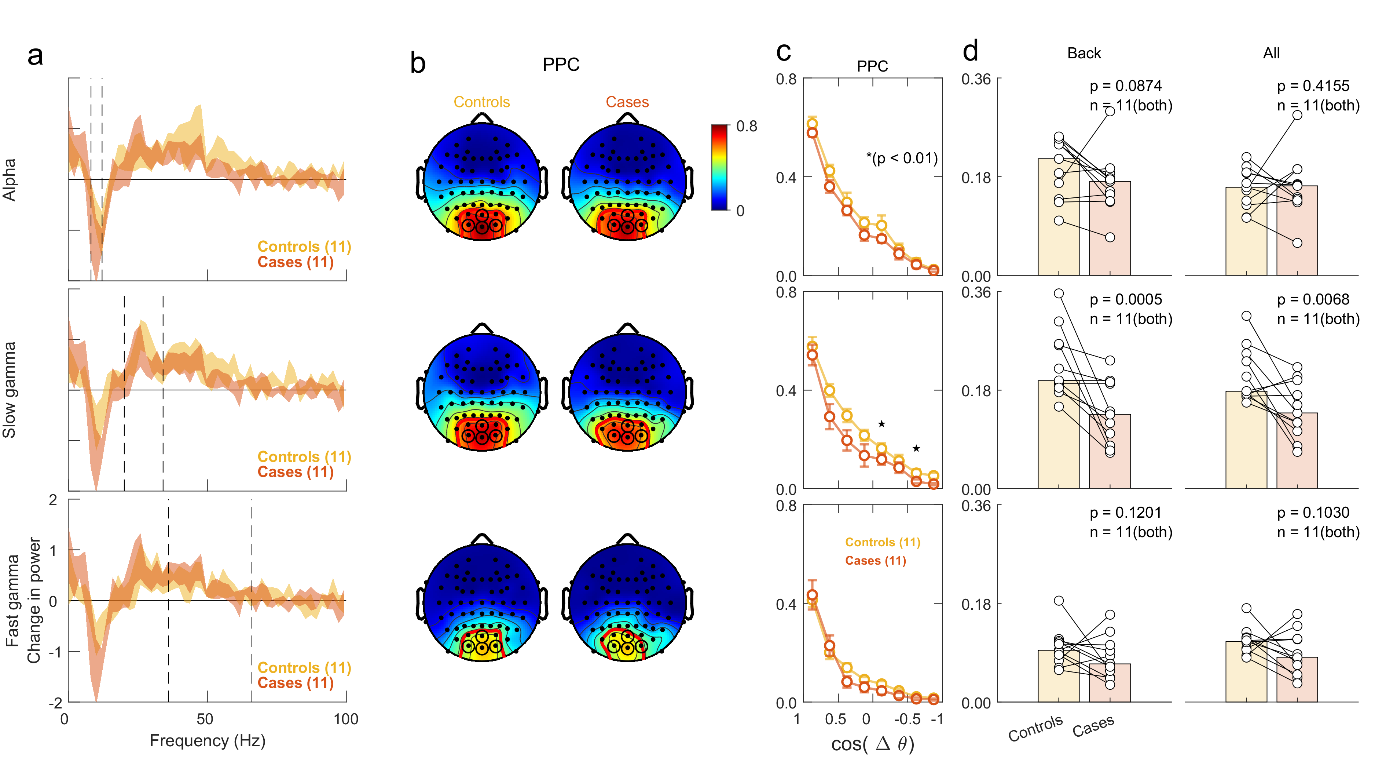

**Figure 5_Supplementary Figure 1**: Similar to Figure 5, with back electrode group as the seed for FC computation.
